# Extended bacterial diversity of the urinary microbiome of reproductive-age healthy European women captured by culturomics and long-read amplicon sequencing

**DOI:** 10.1101/2022.01.19.476882

**Authors:** Svetlana Ugarcina Perovic, Magdalena Ksiezarek, Joana Rocha, Elisabete Alves Cappelli, Márcia Sousa, Teresa Gonçalves Ribeiro, Filipa Grosso, Luísa Peixe

**Author notes:** Correspondence: Luísa Peixe, UCIBIO-REQUIMTE. Laboratory of Microbiology, Faculty of Pharmacy, University of Porto, Portugal. SUP and MK contributed equally to this work.

## Abstract

The recognition of microbiome inhabiting the healthy female bladder engendered the need for comprehensive characterization of the female urinary microbiome (FUM) in health and disease. Although previous studies reported FUM composition at different taxonomic levels, progress towards reliable identification at species level is highly required. The aim of this study was to comprehensively characterize bacterial species of FUM of healthy reproductive-age European women by two complementary methodologies i.e., extended culturomics and long-read third generation sequencing of near full-length 16S rRNA gene.

A wide diversity of bacterial species was captured (297 species) with a median of 53 species/sample, including 16 putative uropathogens. Clustering FUM into community structure types revealed high inter-individual differences. Notably, there was not a single species common to all samples, although the *Lactobacillus* genus was detected in all samples. *Lactobacillus crispatus*, *Lactobacillus iners* and *Lactobacillus mulieris* were observed in high relative abundance in several samples as well as other species (e.g., *Streptococcus agalactiae*, *Atopobium vaginae*, *Gardnerella vaginalis*, *Gardnerella swidsinskii*), while more prevalent species were often low abundant members (e.g., *Finegoldia magna*). We captured remarkable richness within *Corynebacterium* spp. (25 species) and *Lactobacillaceae* (4 genera, 14 species). While amplicon sequencing allowed detection of more anaerobic species (e.g., 11 *Peptoniphilus* spp.), culturomics enabled the identification of recently recognized *Gardnerella* species and putative novel *Corynebacterium* species.

This study provided fine-grained FUM profiling at species level and revealed detailed FUM structure, which is critical to unveil the potential relationship between specific microbiome members and urinary diseases/disorders.

**IMPORTANCE:** Despite evidence of the resident microbial community in the female lower urinary tract, bacterial species diversity and abundance in healthy women is still unclear. This study demonstrated that complementarity between optimized culture-dependent and –independent approaches is highly beneficial for comprehensive FUM species profiling by detecting higher FUM species diversity than previously reported, including identification of unreported *Lactobacillaceae* species and putative novel *Corynebacterium* species. Although some particular species were present in high relative abundance, low-abundant members were more prevalent. FUM classification into community structure types demonstrated high inter-individual differences in urinary microbiome composition among healthy women. We also report moderate correlation between culture-dependent and -independent derived data highlighting drawbacks resulting from each methodological approach. Our findings suggest that FUM bacterial diversity reported from previous studies may be underestimated. Finally, our results contribute to the fundamental knowledge of healthy FUM required for further exploration of the urinary microbiome role in urinary tract diseases.

## Background

Emerging studies in the female urinary microbiome (FUM) have suggested the importance of this unique bacterial community in maintaining urinary tract (UT) healthy (1–6). The advance in FUM characterization through next-generation sequencing and culture-based methodologies has allowed identification of FUM members and indication of their involvement in various UT conditions. These breakthrough findings have triggered the reassessment of current diagnosis practice of urinary tract infection (UTI) (7, 8) and the investigation of still poorly understood etiologies of UT disorders (e.g., overactive bladder syndrome, urgency urinary incontinence and interstitial cystitis/bladder pain syndrome) (9–11).

Up to date, studies have described healthy FUM as a community dominated by certain genera such as *Lactobacillus*, *Gardnerella* or *Streptococcus*, or a mixed community without a single dominant genus involving, e.g., the combination of *Staphylococcus, Corynebacterium* and *Prevotella* genera (10, 12–14). Although the composition of healthy FUM at genus level is relatively established, its species-level composition has not been comprehensively studied. Available studies point to dominance of e.g., *Lactobacillus crispatus, Lactobacillus jensenii*, and *Gardnerella* spp. (often mistakenly reported as *Gardnerella vaginalis*) (15), and the presence of certain potential uropathogens such as *Escherichia coli* and *Enterococcus faecalis,* usually observed in low amounts (7, 13, 16, 17). In fact, detailed species-level characterization is essential to understand FUM diversity and identify key functions contributing to urinary health and disease, since specific features are often species- or even strain-specific.

Current approaches for FUM characterization usually involve culturomics with matrix-assisted laser desorption/ionization time-of-flight mass spectrometry (MALDI-TOF MS) as primary identification method and/or DNA sequencing methodologies targeting individual short hyper-variable regions of the 16S rRNA gene (3, 9, 10, 12, 18). However, some methodological drawbacks concerning the identification at species level of some FUM members can still be recognized. For instance, culture-based methodologies with limited growth conditions and isolates’ identification of insufficient resolution power do not fully capture bacterial species diversity, while short-read DNA-based methods are often limited to a reliable identification of FUM members only at genus level (18).

Considering the above, there is a critical need for accurate and sensitive characterization of the urinary tract microbiome of asymptomatic, healthy individuals to fully support future action in deciphering the microbial community shifts from a eubiotic to dysbiotic state, in order to guide the development of approaches to maintain or restore healthy FUM composition.

To improve our understanding of FUM, we analyzed midstream urine samples of twenty reproductive-age healthy women using complementary approaches of extended culturomics supported by a deep bacterial taxonomic resolution and 16S rRNA long-read sequencing.

## Methods

### Participants and sample collection

This study was approved by the Faculty of Pharmacy (University of Porto, Porto, Portugal) Ethics Committee and written informed consent was obtained from all study participants. Twenty women of reproductive age were recruited between November 2016 and July 2018, following strict criteria: no pregnancy, no symptoms nor diagnosis of current UTI, and no antibiotic exposure in the previous month. A questionnaire was conducted concerning personal and health information that was encrypted, ensuring data confidentiality. Participants were carefully instructed in the collection technique. In the third week of the menstrual cycle, each participant provided a first-morning midstream voided urine sample by self-performed non-invasive procedure via 40 ml sterile containers. Sampling procedure included vaginal swabbing, prior to urine collection, for minimizing cross-contamination.

Urinary dipstick (Combur-Test, Roche) analysis and microscopic examination of the re-suspended sediment of centrifuged urine (1 ml) were performed. Up to 2 hours after collection, urine samples were subjected to extended culturomic protocol, concurrently pre-treated for amplicon sequencing analysis, and stored at -80 °C. The FUM culturomic data from ten women published in the context of urinary tract microbiome temporal stability (17), were included in this study. Since this manuscript includes novel data from amplicon sequencing performed on the same samples, previous culturomic data was used for comparison of efficacy of two methodologies and accurate assessment of community structure types.

### Extended culturomics

The extended culturomic protocol included inoculation of 0.1 ml of urine onto the large plate surface (140 mm diameter) of Columbia agar with 5% sheep blood (blood agar plates - BAPs, Biogerm, Portugal) and HiCrome UTI agar (chromogenic agar plates - CAPs, HiMedia, India) supplemented as previously described (19, 20). BAPs and CAPs were incubated under aerobic and microaerophilic conditions (GENbox MICROAER, bioMérieux, France) at 37 °C for 48 h. Additionally, BAPs were incubated under anaerobic conditions (GENbox ANAER, bioMérieux, France) at 37 °C for 48 h. In case of a suspected high bacterial load based on microscopic observation, ten-fold serial dilutions (up to 0.001) were performed using saline solution (0.9% NaCl) to obtain a countable range of colony forming units (CFU/ml). Each morphologically distinct colony type was counted, and 1-5 colonies of each morphology were further identified. The plate presenting the higher CFU count was considered as the representative count of each isolate in a sample. Relative abundance (RA; %) was calculated by generating the percent of total CFU/sample.

### Identification of cultured bacteria

MALDI-TOF MS with the *in vitro* diagnostic (IVD) database version 3.0 (VITEK MS automation control and Myla software, bioMérieux, France) was used to identify the bacterial isolates. Isolates with no identification, with discrepant results between MALDI-TOF MS identification and phenotypic characteristics, or with known insufficient resolution power for species identification were further subjected to sequencing of 16S rRNA gene and/or other genetic markers (*pheS* for *Lactobacillus* and *Limosilactobacillus*, *cpn60* for *Gardnerella, rpoB* for *Acinetobacter*, *Corynebacterium* or *Staphylococcus,* and *recN* for *Citrobacter*) and/or PCR assays for the detection of species-specific genes (*dltS* for Group B *Streptococcus, sodA* for *Enterococcus faecalis*, and *malB* for *Escherichia coli*) (Supplementary Table S1). GenBank accession numbers and species identification for FUM isolates subjected to Sanger sequencing are available in Supplementary Table S2 and previously published by Ksiezarek *et al*. (17). Phylogenetic analysis based on individual genes were performed to access putative novel species by using MEGA version 7.0 (21), constructed according to neighbour-joining method (22), and genetic distances were estimated using Kimura’s 2-parameter model (23). The reliability of internal branches was assessed from bootstrapping based on 1000 resamplings (24).

### DNA extraction and amplicon sequencing

Samples were pretreated prior to DNA extraction, which included centrifugation of 20 ml of urine at 5,500 rpm for 15 min, and the resulting pellet was suspended in 1 ml of phosphate buffered saline (PBS) and stored at -80 °C until further processing. PBS was discarded by centrifugation at 10,500 rpm/15 min/4°C, immediately before genomic extraction. Genomic DNA from urine samples was extracted using Qiagen DNeasy Blood and Tissue Kit (Qiagen, Germany), according to the manufacturer’s protocol, using pretreatment for Gram-negative bacteria. DNA was eluted into 50 μl of Tris-HCl (pH 8.0) and stored at 4°C. DNA quality was analyzed by agarose gel electrophoresis, and quantity was measured on Qubit dsDNA HS Assay Kit (Invitrogen, Life Technologies, UK). Controls consisting of reagent blanks (washing buffer, lysis buffer and kit reagents) were processed as the urine samples. Since extraction controls showed no traceable amounts of DNA, they were not included for sequencing. PCR amplification of the hypervariable 16S rRNA gene V1-V8 regions sequenced with universal primers (27F:AGAGTTTGATCCTGGCTCAG, and BS-R1407:GACGGGCGGTGWGTRC), library construction and sequencing with SMRT® technology on PacBio RS II sequencing system was provided as a custom service of Eurofins GATC Biotech GmbH (Germany).

### Sequencing data analysis

After sequencing, primers, sequence adaptors, and low base quality calls were removed by Cutadapt. Chimera sequences were checked, and removed by UCHIME (version 4.2.40) (25). The non-chimera and unique sequences were subjected to BLASTn (26) analysis using non-redundant 16S rRNA reference sequences with an E-value cutoff of 1e-06. Reference 16S rRNA gene sequences were obtained from the Ribosomal Database Project Classifier (27). Only good quality and unique 16S rRNA sequences which have a taxonomic assignment were considered and used as a reference database to assign operational taxonomic unit (OTU) status with a 97% similarity. Taxonomic classification was based on the NCBI Taxonomy (28). All the hits to reference 16S rRNA database were considered and specific filters were applied to the hits to remove false positives. The thresholds applied were ≥ 97.00% identity, ≥ 95.00% alignment coverage, 1000 minimum query length, 10% bitscore threshold for multiple hits, and 250 maximum hits to consider for multiples hits. If the final number of high-quality reads after all filtering steps was less than 1000, the corresponding sample was excluded. Finally, RA was calculated by generating the percent of total reads for each sample.

### Statistical analysis

Community structural analyses were done using relative proportions of CFU/ml and reads for each genus and species within individual urine samples. Based on similarity (or dissimilarity) of community composition between samples and taking into account all members and their proportion in a community, we identified community structure types performing hierarchical clustering of Bray-Curtis dissimilarity distance matrices with a cutoff of 0.8, via the package vegan (version 2.5-2) (29) in R (version 3.4.4) (30). Alpha diversity was estimated using Shannon index (H’). Principal coordinates analysis (PCoA) and Mantel test between the dissimilarity distance matrices (based on Bray-Curtis index) were performed to compare structure types obtained by both methodologies. To identify species responsible for community structure differences, a biplot of the PCoA was created using a weighted average of the species scores, based on their RA in the samples. Data visualisation was carried out using gplots (version 3.0.1.1) (31), ggplot2 (version 3.2.1) (32) and eulerr (version 5.1.0) (33) R packages.

## Results

### Overview of the healthy female study cohort

Our study cohort included twenty female participants aged 24-38 years (average = 31; standard deviation = 4). Most women identified themselves as Portuguese nationality (80%) followed by other European nationalities (20%). Average body mass index was 21.9 kg/m^2^. Most women had a normal menstrual cycle (90%) and used contraceptives (85%), with few having experienced at least one pregnancy (25%). Characteristics of our study cohort comprised of healthy highly educated women, including clinical and behavioral questionnaire data (personal medical history, UT health and infection history, pregnancy history, demographic and lifestyle information), and results of urine dipstick and sediment microscopic analysis are available in Supplementary Tables S3 and S4.

### Characterization of community structure types by culturomics

Using extended culturomics we observed a high bacterial load in urine samples (10^3^-10^8^ CFU/ml, 6 10^4^ CFU/ml in 80% of samples). Two thousand and forty-three isolates were studied (median = 103 isolates/sample) and assigned to 131 species (median = 20 species/sample) and 54 genera, as identified either by MALDI-TOF MS and/or sequencing of most suitable genes (Supplementary Table S5). In this cohort, we identified for the first time 13 bacterial species from different genera [*Dermacoccus nishinomiyaensis*, *Gardnerella leopoldii, Gardnerella swidsinskii*, *Gardnerella* genomospecies 3, *Globicatella sulfidifaciens*, *Lactobacillus mulieris*, *Lactobacillus paragasseri*, *Limosilactobacillus urinaemulieris*, *Limosilactobacillus portuensis*, *Limosilactobacillus mucosae* (former *Lactobacillus mucosae*), *Pseudoglutamicibacter cumminsii*, *Staphylococcus carnosus*, and *Staphylococcus equorum*, and 5 putative novel *Corynebacterium* species (Supplementary Table S5 and Fig. S1). Alpha diversity varied from 0.001 to 2.65 (median H′ = 1.5). Bacterial species detected by culturomics and their RA per sample are listed in Supplementary Table S5. Of note, *Corynebacterium* (18 species), *Staphylococcus* (14 species), *Streptococcus* (10 species), *Lactobacillus* (7 species) and *Actinomyces* (6 species) were the genera that presented the highest species-level diversity.

Clustering FUM into community structure types (CST) was performed at genus and species level (samples in the same CST shared >80% similarity by Bray-Curtis distance). Hierarchical clustering at genus level identified 3 CST (Supplementary Fig. S2). The most common CST was CST3 (n=15/20) largely dominated by *Lactobacillus* in combination with other genera (e.g., *Staphylococcus*, *Corynebacterium*, *Streptococcus* and *Cutibacterium*), followed by CST2 (n=4) characterized mostly by *Gardnerella*, and CST1 dominated by *Citrobacter* (n=1). On the other hand, species-level clustering resulted in 13 CST (Fig. 1, Table 1), mostly representing individual urine specimens as only 5 CST included more than one sample. With exception of 2 clusters dominated by a single bacterial species (CST1-*Citrobacter koseri*, and CST2-*Gardnerella vaginalis*, >90%), the remaining CST were predominantly represented by an extraordinarily diverse bacterial community (different combinations and RA of bacterial species), which varied widely from 1.21±0.05 to 2.65 as calculated by the Shannon diversity index (Fig. 1, Table 1). For instance, CST5 was characterized by combination of *Lactobacillus iners* with other bacterial species, CST12 included *Lactobacillus crispatus, Lactobacillus mulieris* and other bacterial species, while CST10 comprised abundant *Atopobium vaginae*, low abundant *Streptococcus anginosus* and in one sample highly abundant *Gardnerella swidsinskii* (RA ∼ 50%) (Fig. 1).

**Figure 1.**
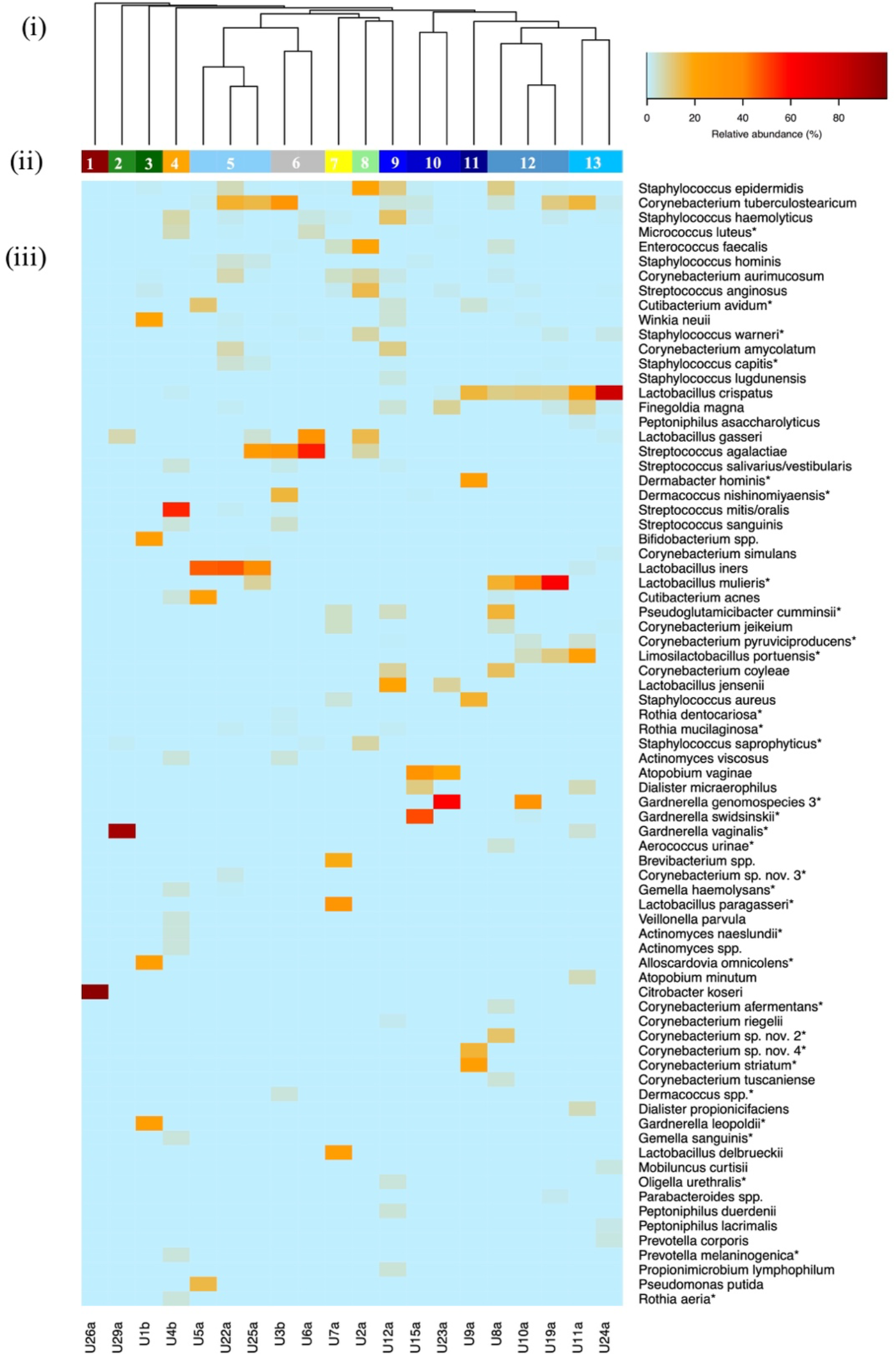
Species-level community structure types of healthy FUM by culturomics. (i) Hierarchical clustering of Bray-Curtis dissimilarity distance matrices on the relative proportions of CFU/ml within individual urine samples. (ii) Bars below dendrogram denote community structure types. (iii) Heatmap of RA of bacterial species within each urinary microbiome. Only species that are at least 1% abundant in at least one sample are shown in order of decreasing prevalence (from top to bottom). Asterisk denotes detection only by culturomics and not by amplicon sequencing.

**Table 1.** Overview of all healthy community structure types and their characteristic species by culturomics. Shared species within a structure type are presented in order of decreasing relative abundance (relative abundance > 1%, only top 5 shown).

| Structure type | Characteristic species | Samples | Shannon index<br>(mean $H'$ ± Standard Deviation) |
| --- | --- | --- | --- |
| 1 | *Citrobacter koseri* | U26a | 0.001 |
| 2 | *Gardnerella vaginalis* | U29a | 0.33 |
|  | *Lactobacillus gasseri* |  |  |
| 3 | *Gardnerella leopoldii*<br>*Alloscardovia omnicolens*<br>*Bifidobacterium spp.*<br>*Winkia neuii*<br>*Streptococcus anginosus* | U1b | 1.61 |
| 4 | *Streptococcus mitis/oralis*<br>*Staphylococcus haemolyticus*<br>*Micrococcus luteus*<br>*Actinomyces spp.*<br>*Lactobacillus crispatus* | U4b | 1.85 |
| 5 | *Lactobacillus iners*<br>*Corynebacterium tuberculostrictum*<br>*Staphylococcus epidermidis*<br>*Staphylococcus hominis*<br>*Staphylococcus capitis* | U5a,<br>U22a,<br>U25a | 1.61±0.20 |
| 6 | *Streptococcus agalactiae*<br>*Streptococcus salivarius/vestibularis*<br>*Micrococcus luteus*<br>*Staphylococcus haemolyticus* | U3b,<br>U6a | 1.40±0.35 |
| 7 | *Lactobacillus paragasseri*<br>*Lactobacillus delbrueckii*<br>*Brevibacterium spp.*<br>*Pseudoglutamicibacter cummingsii*<br>*Corynebacterium jeikeium* | U7a | 1.87 |
| 8 | *Enterococcus faecalis*<br>*Staphylococcus epidermidis*<br>*Lactobacillus gasseri*<br>*Streptococcus anginosus*<br>*Corynebacterium aurimucosum* | U2a | 1.97 |
| 9 | *Lactobacillus jensenii*<br>*Staphylococcus haemolyticus*<br>*Staphylococcus epidermidis*<br>*Corynebacterium amycolatum*<br>*Corynebacterium coyleae* | U12a | 2.65 |
| 10 | *Atopobium vaginae*<br>*Streptococcus anginosus* | U15a,<br>U23a | 1.21±0.05 |
| 11 | *Corynebacterium striatum*<br>*Dermabacter hominis*<br>*Staphylococcus aureus*<br>*Corynebacterium* sp. nov. 4<br>*Lactobacillus crispatus* | U9a | 1.72 |
| 12 | *Lactobacillus crispatus*<br>*Lactobacillus mulieris*<br>*Staphylococcus epidermidis*<br>*Cutibacterium avidum* | U8a,<br>U10a,<br>U19a | 1.77±0.45 |
| 13 | *Lactobacillus crispatus*<br>*Corynebacterium tuberculostearicum*<br>*Finegoldia magna* | U11a,<br>U24a | 1.52±0.58 |

### Characterization of community structure types by amplicon sequencing

A total of 58,534 reads were generated, with most of them being assigned to the species level (88%; 51,317 reads). One sample (U6a) had <1000 reads and was excluded from the analysis, while for the remaining a median of 2493 reads/sample (interquartile range, IQR 1625 - 3920) was generated. A total of 231 species (IQR 5-115, median = 39 species/sample) belonging to 107 genera and 8 phyla were identified. The alpha diversity varied from 0.135 to 2.79 (median H′ = 0.90). Bacterial species detected by amplicon sequencing and their RA are listed in Supplementary Table S6. Of note, *Corynebacterium* (16 species), *Peptoniphilus* (11 species), *Anaerococcus* (10 species), *Streptococcus* (9 species) and *Bacteroides* (8 species) were the genera that presented the highest species-level diversity.

The same FUM clustering approach was applied to amplicon sequencing data. Genus-level clustering resulted in 5 CST (Supplementary Fig. S3). The *Lactobacillus* genus in combination with other bacterial genera (e.g., *Prevotella, Dialister,* and *Corynebacterium*) represented the most prevalent CST (CST5; 79%, n=15/19). Species-level clustering resulted in 7 CSTs (Fig. 2, Table 2), with the 3 (n=15) most common (being characterized by combination of a highly abundant *Lactobacillus* species (CST3-*L. iners*, CST5-*Lactobacillus gasseri*, CST7-*L. crispatus*) and species from other genera (Fig. 2). Remarkably, the *Lactobacillus iners* enriched-CST was characterized by a reduced species diversity (CST3, H′ = 0.56±0.42) compared to other *Lactobacillus* CSTs. The remaining CSTs included highly abundant *C. koseri* (CST1; n=1/19), *Atopobium vaginae* (CST2; n=1/19), or combination of different species (CST4: *Anaerococcus tetradius* and *Prevotella timonensis*; CST6: *Ralstonia mannitolilytica* and *Streptococcus agalactiae*; n=1 each).

**Figure 2.**
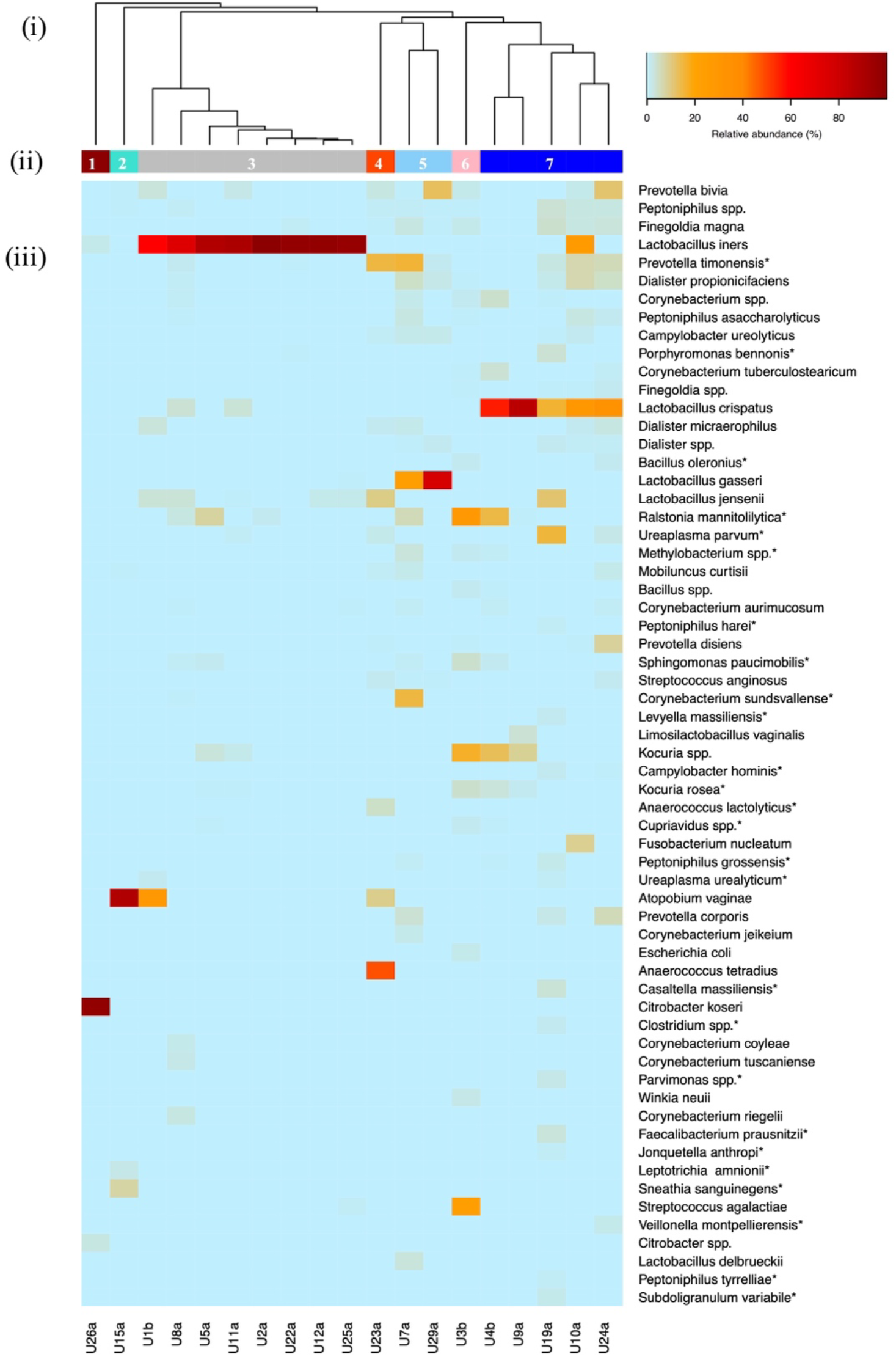
Species-level community structure types of healthy FUM by amplicon sequencing. (i) Hierarchical clustering of Bray-Curtis dissimilarity distance matrices on the relative proportions of reads for each OTU within individual urine samples. (ii) Bars below dendrogram denote community structure types. (iii) Heatmap of RA of bacterial species within each urinary microbiome. Only species that are at least 1% abundant in at least one sample are shown in order of decreasing prevalence (from top to bottom). Asterisk denotes detection only by amplicon sequencing and not by culturomics.

**Table 2.** Overview of all healthy community structure types and their characteristic species by amplicon sequencing. Shared species within a structure type are presented in order of decreasing relative abundance (relative abundance > 1%, only top 5 shown).

| **Structure type** | **Characteristic species** | **Sample s** | **Shannon index**<br>(mean H'± Standard Deviation) |
| --- | --- | --- | --- |
| 1 | *Citrobacter koseri* | U26a | 0.21 |
|  | *Citrobacter* spp.<br>*Lactobacillus iners* |  |  |
| 2 | *Atopobium vaginae*<br>*Sneathia sanguinegens* | U15a | 0.59 |
| 3 | *Lactobacillus iners*<br>*Prevotella timonensis* | U1b,<br>U8a,<br>U5a,<br>U11a,<br>U2a,<br>U22a,<br>U12a,<br>U25a | 0.56±0.42 |
| 4 | *Anaerococcus tetradius*<br>*Prevotella timonensis*<br>*Lactobacillus jensenii*<br>*Atopobium vaginae*<br>*Ureaplasma parvum* | U23a | 1.78 |
| 5 | *Lactobacillus gasseri*<br>*Prevotella timonensis*<br>*Dialister propionificiens*<br>*Campylobacter ureolyticus* | U7a,<br>U29a | 1.70±1.13 |
| 6 | *Ralstonia mannitolilytica*<br>*Streptococcus agalactiae*<br>*Kocuria* spp. | U3b | 2.12 |
| 7 | *Lactobacillus crispatus*<br>*Corynebacterium* spp. | U4b,<br>U9a,<br>U19a, | 1.86±0.85 |
|  | *Corynebacterium*<br>*tuberculostrictum*<br>*Peptoniphilus* spp. | U10a,<br>U24a |  |

### Correlation between community structure types assigned by culturomics and amplicon sequencing

A moderate correlation was observed using the Mantel test (r = 0.5, p< 0.05) between the CST assigned by culturomics and amplicon sequencing. Congruence was observed for the types of highly abundant *C. koseri* and combinations of different *Lactobacillus* species (Fig. 1, Fig. 2), with 37% of samples (7/19) clustering into the same CST by both methodologies and additional 26% of samples (5/19) clustering in the proximate CST (Supplementary Fig. S4).

*Lactobacillus* amongst others were responsible for the reduction in correlation between CST detected by different methodologies (e.g., *Lactobacillus iners* was more frequently detected in a higher RA by amplicon sequencing, while *Cutibacterium acnes* was more frequently detected by culturomics) (Fig. 3). Overall, amplicon sequencing enabled the detection of bacteria difficult to grow by conventional methods (e.g., *Ureaplasma urealyticum, Ureaplasma parvum*) and improved detection of fastidious bacterial species (e.g., *Campylobacter ureolyticus*, *Finegoldia magna, Atopobium vaginae*), whereas culturomics allowed the identification of various Gram-positive bacteria (e.g., *Enterococcus faecalis, Streptococcus agalactiae, Streptococcus anginosus*). Remarkably, some species detected by sequencing in low-reads count (e.g., *Staphylococcus aureus* and *Actinomyces urogenitalis*, RA < 0.1%) were identified by extended culturomics, confirming their presence in a given sample (Supplementary Tables S5 and S6). Culturomics also allowed precise identification of closely related and/or newly described bacterial species e.g., *Gardnerella leopoldii, Gardnerella* genomospecies 3, *Gardnerella swidsinskii, Limosilactobacillus portuensis, Limosilactobacillus urinaemulieris, Lactobacillus paragasseri, Lactobacillus mulieris*, and putative novel *Corynebacterium* species.

**Figure 3.**
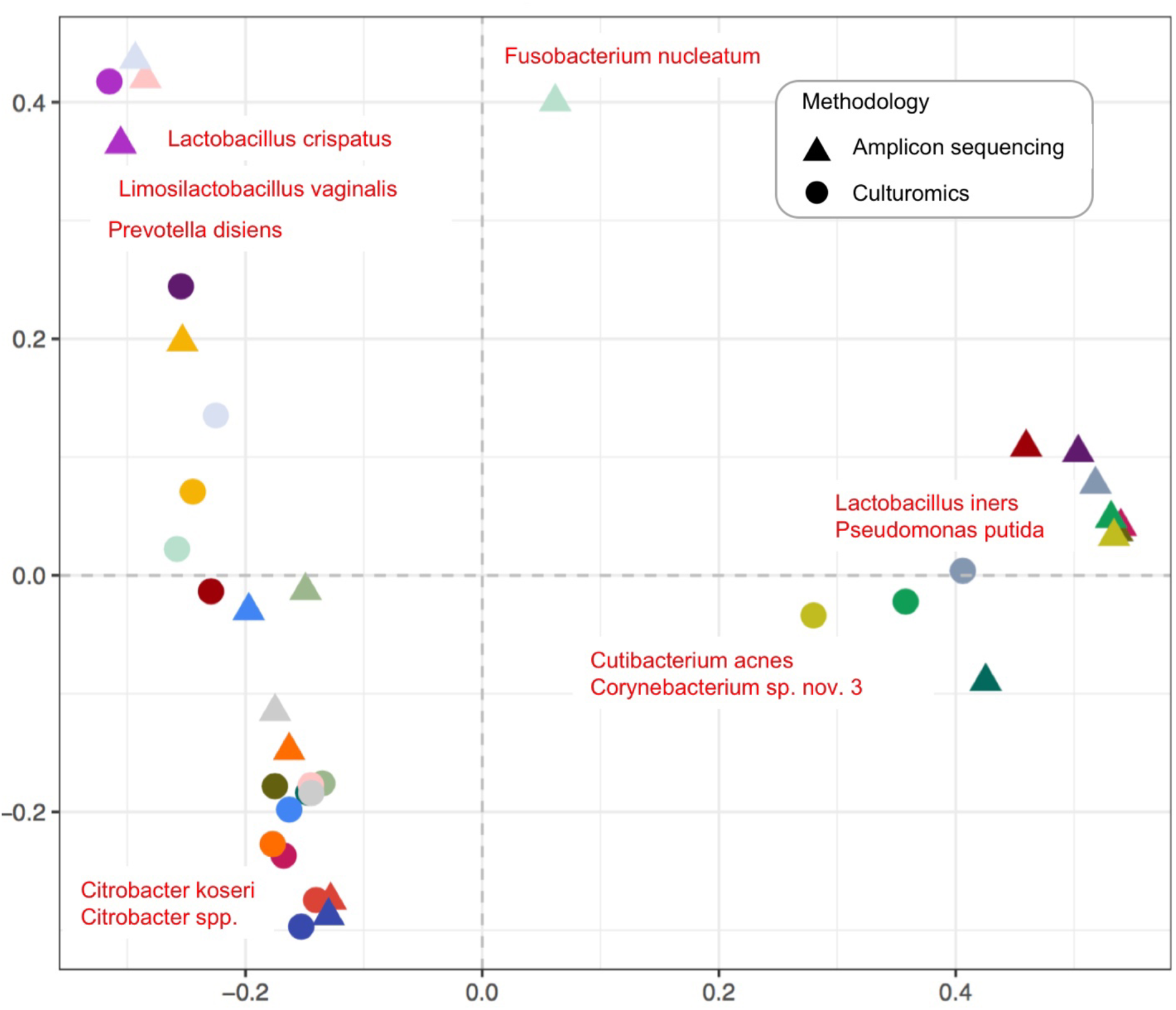
Biplot of the principal coordinate analysis (PCoA) based on the species-level Bray-Curtis dissimilarity matrices. Two-dimensional distances identify dissimilarities between bacterial community structures detected by culturomics and amplicon sequencing. The biplot, based on weighted average of the species scores, shows the top 10 species with the largest contributions to dissimilarities. Same colour indicates the same sample.

### Overview of bacterial species in healthy FUM

In total we captured an extended set of bacteria belonging to 8 phyla, 116 genera and 297 species (median = 53 species/sample) in healthy FUM (Supplementary Tables S5 and S6; Supplementary Fig. S5). Out of 297 species, we have identified 65 species (22% of total species) belonging to 35 genera and 5 phyla by both methodologies. Certain genera were characterized by outstandingly high species-level diversity that could be captured only by combined culture-based and DNA-dependent approaches. For instance, from a total of 25 *Corynebacterium* species, 8 could be identified by both methodologies (apart from 10 detected only by extended culturomics – including 5 putative novel species – and 7 only by amplicon sequencing), or from 14 species belonging to *Lactobacillaceae* (4 genera), 7 could be detected by both methodologies (in addition to 4 identified only by extended culturomics and 3 by amplicon sequencing) (Supplementary Tables S5 and S6).

We could not identify a single species present in all samples, although the genus *Lactobacillus* was detected in all. Instead, we were able to unveil 14 prevalent bacterial species (present in more than 50% of samples) with at least 1% of abundance in one sample (Fig. 4, Supplementary Table S7). *Staphylococcus epidermidis* was the most common species (n=18/20), followed by *Finegoldia magna* (n=16/20), *Corynebacterium tuberculostearicum* (n=15/20), and *Prevotella bivia* (n=15/20) (Supplementary Table S7). Remarkably, the common species were mostly low-abundant members (RA < 5%) (Fig. 4).

**Figure 4.**
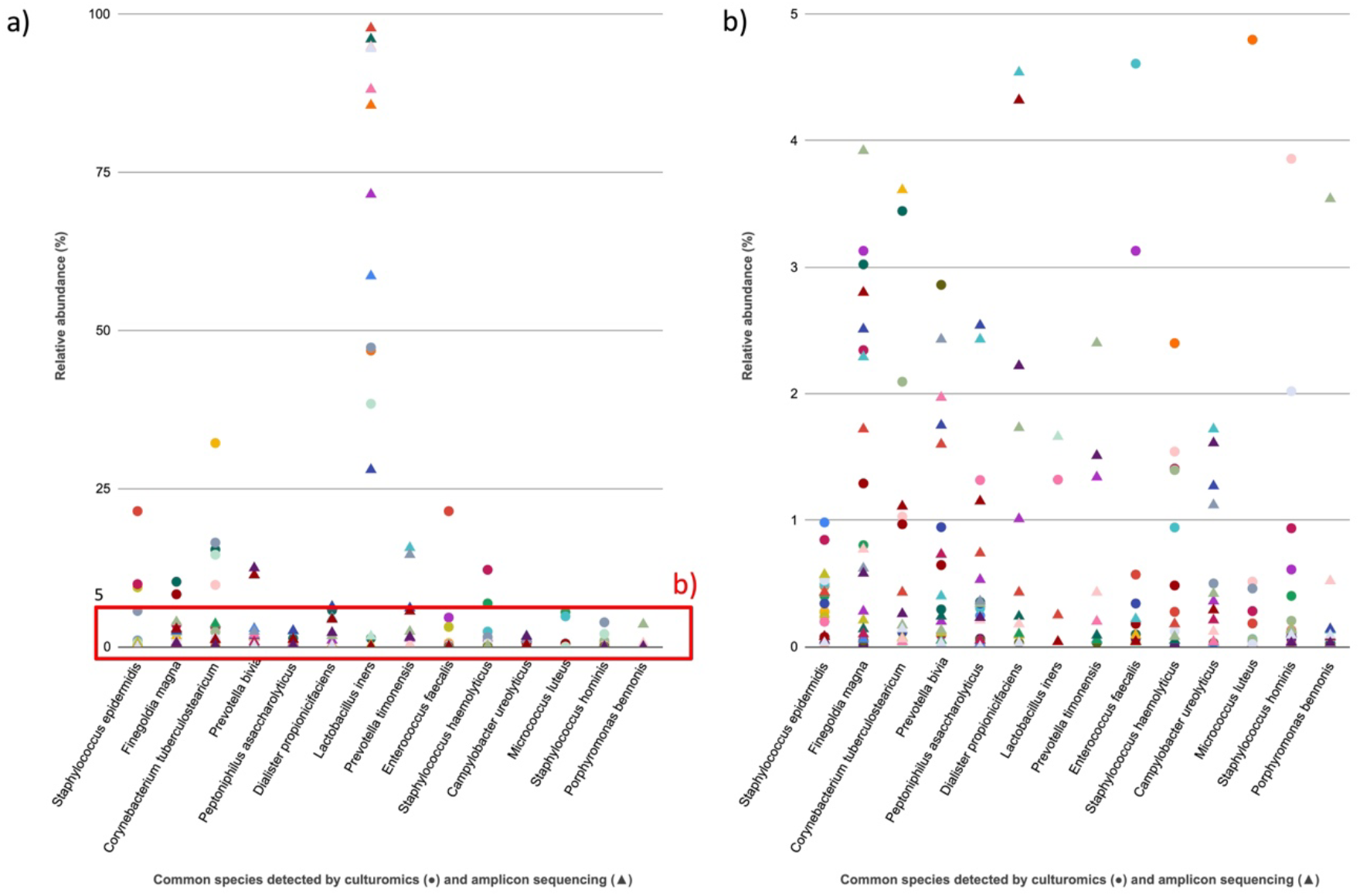
Common bacterial species of healthy FUM detected by culturomics and amplicon sequencing. (a) RA per sample of species present in more than 50% of samples by culturomics and amplicon sequencing. Only species that are detected by culturomics or amplicon sequencing with at least 1% abundance in at least one sample are presented in order of decreasing prevalence (from left to right). Same colour indicates the same sample. (b) Close-up of section of Fig. 4(a) showing the RA range 0.01-5%.

Additionally, we looked for the presence of opportunistic pathogens associated with the healthy urogenital tract and found 16 bacterial species largely varying in their RAs (IRQ 0,03-96.62%), among which *Enterococcus faecalis*, *Streptococcus anginosus*, and *Ureaplasma parvum* were the most frequently identified by both methodologies (Table 3). Noteworthy, *C. koseri* was a highly abundant member detected by both methodologies, while *Atopobium vaginae* was only detected by amplicon sequencing. All opportunistic pathogens associated with the urogenital tract detected by culturomics and/or amplicon sequencing in healthy FUM are listed in Table 3.

**Table 3.** Opportunistic pathogens associated with the urogenital tract. Species are listed in order of decreasing detection frequency in FUM.

| Species | Frequency<br>in FUM*<br>(%) | Culturomics |  | Amplicon sequencing |  |
| --- | --- | --- | --- | --- | --- |
|  |  | Frequency in<br>20 samples<br>(%) | RA<br>(%) | Frequency in<br>19 samples (%) | RA<br>(%) |
| *Enterococcus faecalis* | 12/20<br>(60%) | 11/20<br>(55%) | 0.01-20 | 3/19<br>(16%) | 0.04-0.22 |
| *Streptococcus*<br>*anginosus* | 11/20<br>(55%) | 10/20<br>(50%) | 0.06-13.33 | 7/19<br>(37%) | 0.03-1.31 |
| *Ureaplasma parvum* | 8/20<br>(40%) | ND | ND | 8/19<br>(42%) | 0.10-15.10 |
| *Escherichia coli* | 6/20<br>(30%) | 4/20<br>(20%) | 0.02-0.28 | 4/19<br>(21%) | 0.04-1.54 |
| *Streptococcus*<br>*agalactiae* | 6/20<br>(30%) | 6/20<br>(30%) | 0.03-55.96 | 2/19<br>(10%) | 0.85-23.20 |
| *Ureaplasma*<br>*urealyticum* | 5/20<br>(25%) | ND | ND | 5/19<br>(26%) | 0.08-1.38 |
| *Atopobium vaginae* | 4/20<br>(20%) | 2/20<br>(10%) | 21.66-36.44 | 4/19<br>(21%) | 0.09-86.63 |
| *Staphylococcus*<br>*aureus* | 3/20<br>(15%) | 3/20<br>(15%) | 0.33-16.68 | 1/19<br>(5%) | 0.04 |
| *Staphylococcus saprophyticus* | 3/20<br>(15%) | 3/20<br>(15%) | 0.79-6.67 | ND | ND |
| *Corynebacterium coyleae* | 3/20<br>(15%) | 3/20<br>(15%) | 0.08-12.70 | 3/19<br>(16%) | 0.09-1.54 |
| *Citrobacter koseri* | 3/20<br>(15%) | 1/20<br>(5%) | 99.98 | 3/19<br>(16%) | 0.03-96.62 |
| *Actinotignum schaalii* | 2/20<br>(10%) | 1/20<br>(5%) | 0.28 | 2/19<br>(10%) | 0.08-0.55 |
| *Aerococcus urinae* | 2/20<br>(10%) | 2/20<br>(10%) | 0.12-3.17 | ND | ND |
| *Alloscardovia omnicolens* | 1/20<br>(5%) | 1/20<br>(5%) | 24.55 | ND | ND |
| *Pseudomonas putida* | 1/20<br>(5%) | 1/20<br>(5%) | 14.72 | ND | ND |
| *Stenotrophomonas maltophilia* | 1/20<br>(5%) | 1/20<br>(5%) | 0.13 | ND | ND |

\*Total detection in FUM of 20 participants by both methodologies.
ND - not detected; RA - relative abundance

## Discussion

Understanding the microbial composition of the lower urinary tract in healthy individuals is essential so that microbial changes associated with urinary disorders can be recognized and modulated as a therapeutic strategy. In this study, using a complementary approach supported by two comprehensive methodologies (extended culturomics and amplicon sequencing), we expanded the knowledge on compositional bacterial species patterns of the female lower urinary tract microbiome.

Each technique presented a different capacity to characterize microbiome profiles (∼63% of CST overlap for both methodologies), and only 22% of bacterial species were detected by both methodologies. Predictably, amplicon sequencing allowed more frequent detection of slow-growing species (e.g., *Campylobacter ureolyticus*), and obligate anaerobes (e.g., *Finegoldia magna*) that require particular culturing conditions. Interestingly, amplicon sequencing also revealed high species diversity within certain anaerobic genera (e.g., *Anaerococcus, Peptoniphilus*), however it is unclear if all these species were viable at the time of detection. On the other hand, the cultured isolates could be accurately identified to the species and strain level, thus providing a higher level of resolution, and allowing further investigation to unveil their symbiotic or pathogenic potential. Moreover, some species detected in low-reads count were also identified by extended culturomics, which supports that FUM bacterial diversity reported from DNA-based studies may be underestimated, as also pointed out by other studies (34). Overall, the complementarity of both methodological approaches allowed for a more comprehensive description of the FUM diversity.

Clustering FUM at genus level revealed that the most prevalent CST was characterized by the combination of highly abundant *Lactobacillus* and other genera, confirming previously reported high occurrence of *Lactobacillus* in the urinary microbiome (3, 10, 13, 17). At species level the diversity largely increased, with the majority of the CST being represented by different *Lactobacillus* or *Gardnerella* species in different RA, and in combination with species from other genera, including low-abundant FUM members, as observed in our previous study (17).

Remarkably, we identified for the first time a CST dominated by *Atopobium vaginae* (RA 33-87%) in an asymptomatic individual (U15a) (Fig. 2), in combination with *Gardnerella swidsinskii* (RA ∼ 50%) (Fig. 1) (Supplementary Tables S5 and S6). Although *Atopobium vaginae* is associated with bacterial vaginosis (35, 36), this woman did not report any symptoms associated with urogenital diseases. Potentially, presence of *Atopobium vaginae*, opportunistic uropathogens (e.g., *E. coli*, *C. koseri*, or *E. faecalis*), or species more frequently isolated from women with specific urinary disorders (e.g., *Aerococcus urinae*, *Lactobacillus gasseri*) (Table 3, Fig. 1) (37, 38), might not be sufficient biomarkers of urogenital infections/disorders, since there are high functional variations at strain level. Further elucidation of the function of urinary microbiome members, including characterization (presence and expression) of virulence factors *sensu stricto* playing a significant role in pathogenesis, will likely help to understand the development of urogenital diseases (39, 40).

Interestingly, we detected an outstandingly high diversity of *Corynebacterium* species and *Lactobacillaceae* members that was never reported in previous studies characterizing the asymptomatic FUM (Supplementary Tables S5 and S6) (1–3, 7, 10, 13, 41–43). We also identified 4 *Gardnerella* species in healthy FUM, according to recent genus reclassification (15). This demonstrates that the high number of colonies studied and reliable identification of isolates by specific genotypic markers, together with the usage of cutting-edge long-read third generation sequencing of the 16S rRNA gene increase the knowledge on the composition of bacterial community to the species level in microbiome studies (17, 18, 44).

Additional strengths of this study include sample processing up to 2 hours after collection, allowing us to identify anaerobic bacteria that seems to significantly contribute to the urinary microbiome repertoire (34), but are rarely or not reported by other healthy FUM culturomics studies (e.g., *Prevotella corporis)* (3, 7, 45). Methodological improvements include also the use of a larger volume sample size (20 ml), compared to previously used urine volume (mostly 1 ml) in DNA extraction protocols, which increased high-quality microbial DNA yield required for high-resolution sequencing, and unveiled detection of species not previously reported in DNA-based studies (e.g., *Alistipes putredinis*) (1, 10, 12, 34). Another important improvement was the use of a cutting-edge sequencing technique, including near full-length 16S rRNA gene sequencing using PacBio SMRT cell technology (18, 46–48), and appropriate gene markers to identify cultured isolates at species level, which enable increased taxonomic resolution, as well as validation of several low-read sequencing data (< 0.1% RA) by our extended culturomic protocol.

One limitation of this study was the small cohort size, yet our strictly selected participants (e.g., no antibiotics for any medical reason within the month prior to urine collection and samples collected on 3^rd^ week of menstrual cycle) represented a homogeneous healthy female group. Another limitation of this study could be the use of voided urine instead of urine collected by suprapubic aspiration or urethral catheterization (49, 50). However, suprapubic aspiration or catheterization of participants who were not at a high risk of bacterial infection or without any clinical urinary symptoms was not ethically feasible as per our local ethics committee and, in fact, voided urine is a sample commonly used for diagnosis of urinary tract pathologies. Additionally, voided urine samples capture the urethral bacteria which can play an important role in urinary tract conditions. Moreover, in our study, careful vaginal swabbing was employed right before urine collection, for i) minimizing vulvo-vaginal bacterial contribution; ii) assessing similarity of paired urinary tract and vaginal microbiome, which according to our preliminary data, is substantially different for most women (unpublished data).

## Conclusions

Our study substantially enlarged the knowledge on bacterial species diversity in healthy FUM and provided extensive taxonomic characterization of *Gardnerella, Lactobacillaceae*, and *Corynebacterium* which are prevalent members in this niche. We demonstrated that, at species level, healthy FUM is highly diverse within and between individuals, and the most prevalent FUM members are low-abundant bacteria, potentially playing an important role in urinary tract eubiosis.

This study provides a fine-grained analysis using improved culture- and DNA-based approaches that were shown to be highly beneficial to capture FUM species-level diversity. Additionally, the data provided here can be useful to estimate the bias resulting from using just one methodology.

Finally, our findings provide essential species level information for further studies on microbiome dysbiosis associated with urinary tract infection and lower urinary tract symptoms, required for development of more effective diagnostic and/or therapeutic strategies. As we begin to detect near full composition and diversity of the urinary microbiome, future studies accessing the functionality of the resident microbiome in the human urinary tract should receive high priority.

## List of abbreviations

BAP: Columbia agar with 5% sheep blood plate

CAP: chromogenic agar plate

CFU: colony forming unit

CST: community structure type

*dltS*: histidine kinase specific to group B *Streptococcus*

FUM: female urinary microbiome

IVD: *in vitro* diagnostic

*malB*: maltose operon protein B

MALDI-TOF MS: matrix-assisted laser desorption/ionization time-of-flight mass spectrometry

ND: not detected

OTU: operational taxonomic unit

PCoA: principal coordinates analysis

*pheS*: phenylalanyl-tRNA synthetase alpha subunit

RA: relative abundance

*recN*: DNA repair protein

*rpoB*: RNA polymerase beta subunit

SDS: sodium dodecyl sulfate

*sodA*: superoxide dismutase

UT: urinary tract

UTI: urinary tract infection

## Declarations

### Ethics approval and consent to participate

Approval of the study was obtained from the Faculty of Pharmacy (University of Porto, Porto, Portugal) Ethics Committee. Procedures performed in the study were all in accordance with the ethical standards of the institutional and national research committee, with the 1964 Helsinki Declaration, and its later amendments. All individual participants included in the study had given written informed consent.

### Consent for publication

Not applicable.

### Availability of data and materials

The datasets supporting the conclusions of this article are included within the article and its supplementary material, and available in the Sequence Read Archive repository, under BioProject accession number PRJNA548360 (51).

## Authors’ contributions

SUP, MK, FG and LP designed the study and supervised participant recruitment. SUP, MK and JR processed the samples and collected the data. SUP, MK, JR, MS, EAC and TGR performed the isolates’ identification. TGR supervised MS and EAC. SUP conducted the community data analysis and visualization. SUP and MK interpreted the data and wrote the manuscript. TGR, FG and LP revised the article. All authors read and approved the manuscript.

## Acknowledgements

We thank the women who participated and contributed their time and effort to make this study a success. We thank Helena Ramos and Paulo Pinto (Hospital Geral de Santo António, Porto, Portugal) for their technical support with MALDI-TOF MS-based microbial identification. The bioMérieux (Portugal, Lda.) provided equipment and material for MALDI-TOF MS analysis, and had no role in the study design, data collection and analysis, decision to publish, or writing of the manuscript.

## Supplementary Tables S1-S7

https://docs.google.com/spreadsheets/d/1ctw656zxecl2TVC2Kr3bvvdM3boW1MCu/edit

Table S1. PCR primers used for bacterial species identification

Table S2. Accession numbers and species identification for FUM isolates subjected to Sanger sequencing

Table S3. Demographics, health data of the study participants (N = 20)

Table S4. Results of urine dipstick and sediment microscopic analysis

Table S5. Species detected by culturomics and their RA in all samples

Table S6. Species detected by amplicon sequencing and their RA in all samples

Table S7. Species detected by culturomics and amplicon sequencing in more than 50% of samples and their RA.

---

**Figure S1.**
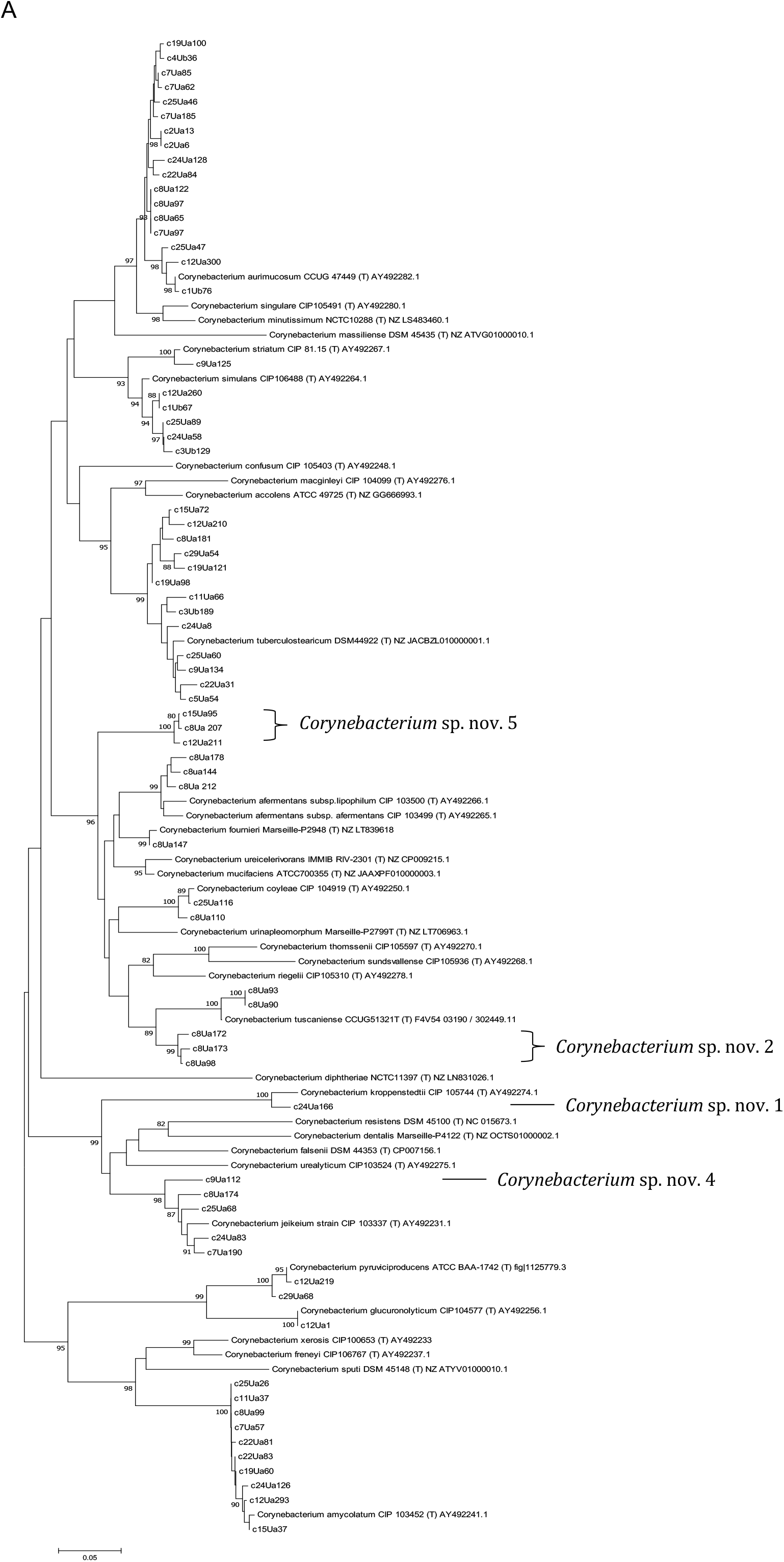
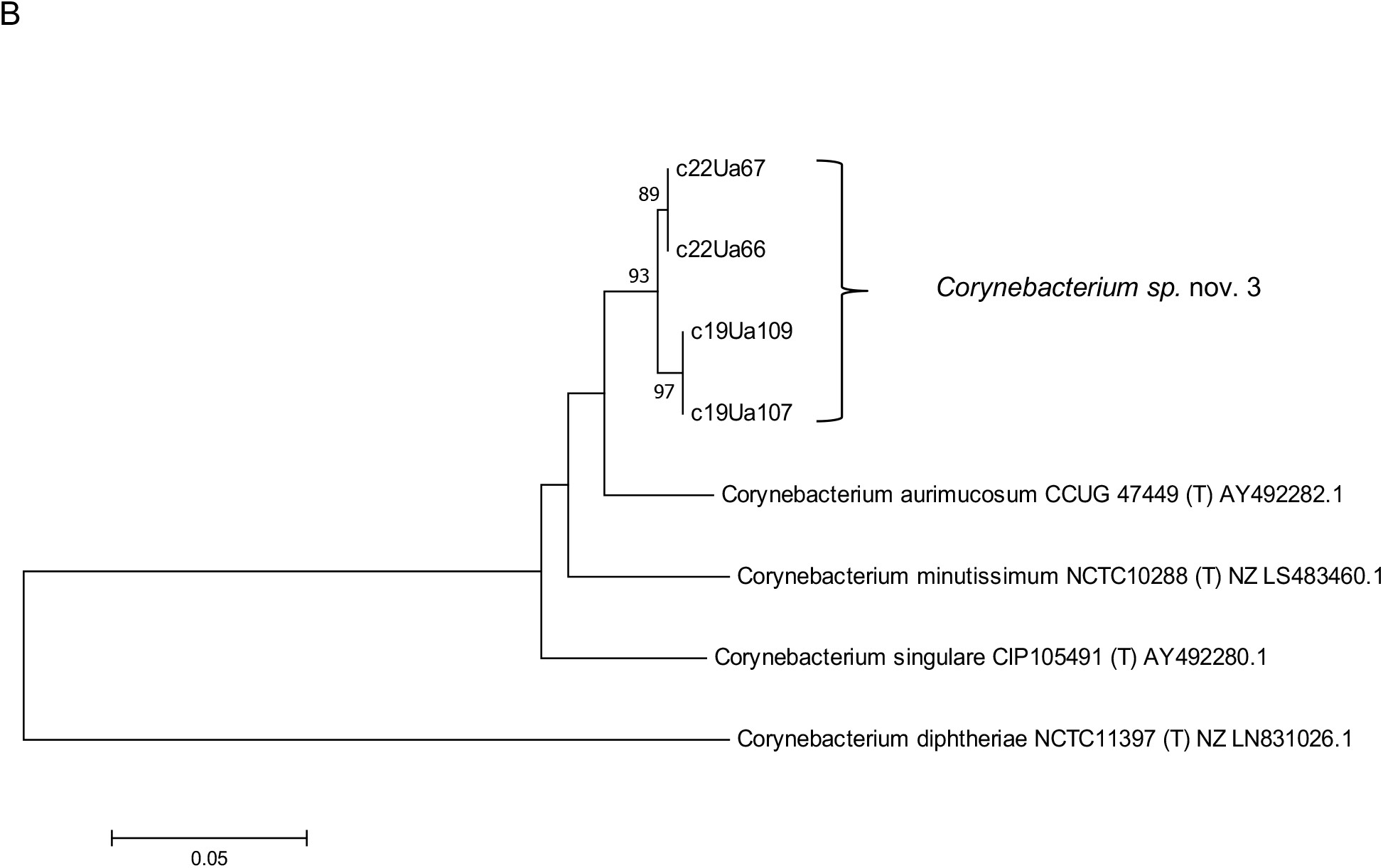
*Corynebacterium* putative novel species 1-5. Neighbor-joining tree based on *rpoB* gene sequences showing the phylogenetic relationships between *Corynebacterium* selected closely related type strains and putative novel species. Nucleotide sequences were extracted from draft/complete genomes obtained from the NCBI Assembly Database, for which the accession numbers are shown next to the strain designation. Bootstrap percentages (based on 1000 replications) are shown at nodes. Only values above 80% are shown. Bar, 0.05 substitutions per nucleotide position. Figure S1.A represents isolates sequenced with primers F- CGWATGAACATYGGBCAGGT and R- TCCATYTCRCCRAARCGCTG with ∼450 bp amplicon, while figure S1.B includes isolated sequenced with primers F- CNTCBCACTAYGGNCGNATG and R- GAVCGNGCGTGRATCTTYTC with ∼1700 bp amplicon.

**Figure S2.**
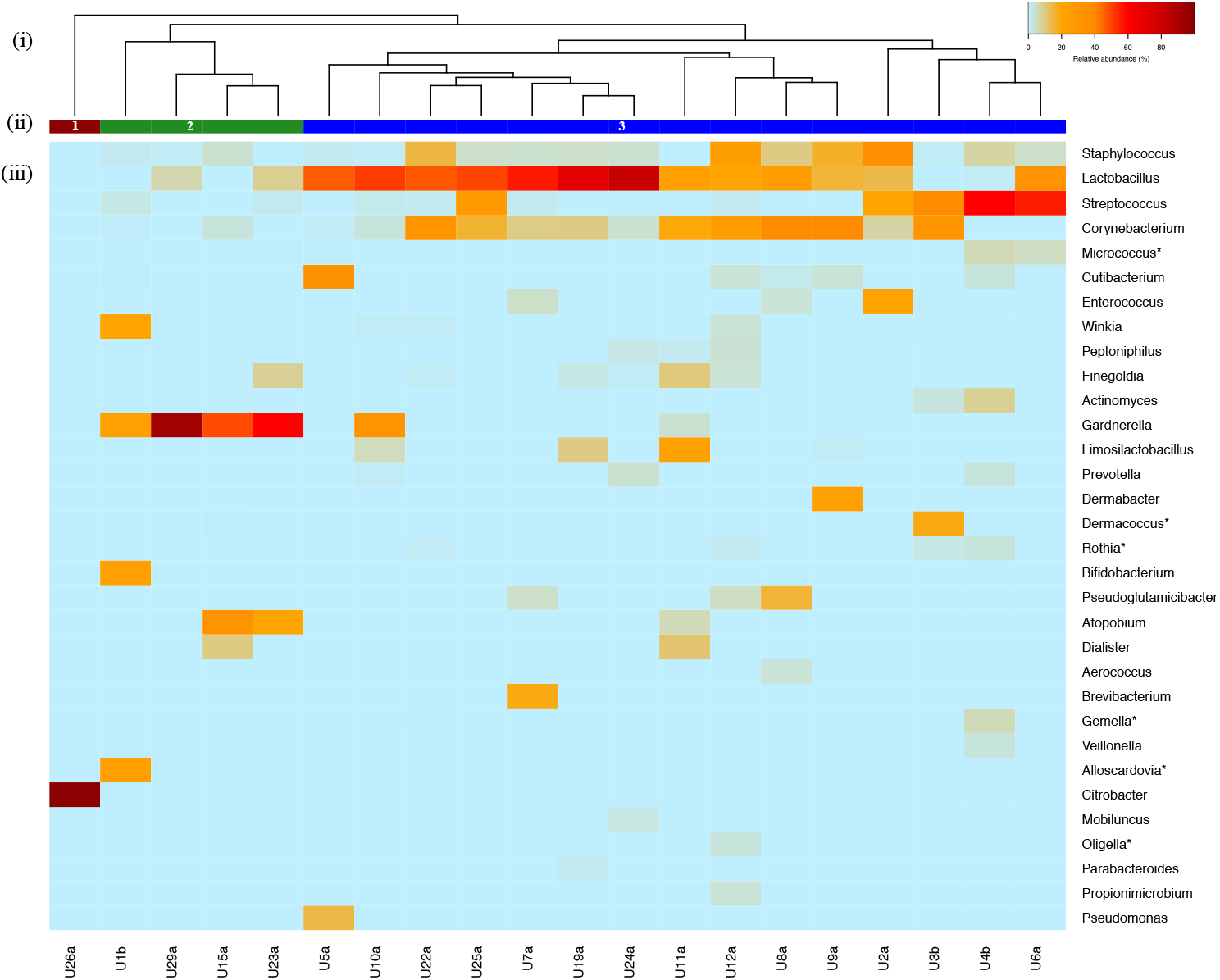
Genus-level community structure types of healthy FUM by culturomics. (i) Hierarchical clustering of Bray-Curtis dissimilarity distance matrices on the relative proportions of CFU/ml within individual urine samples. (ii) Bars below dendrogram denote community structure types. (iii) Heatmap of relative abundances of bacterial genera within each urinary microbiota. Only genera that are at least 1% abundant in at least one sample are shown in order of decreasing prevalence (from top to bottom). Asterisk denotes detection only by culturomics and not by community amplicon sequencing.

**Figure S3.**
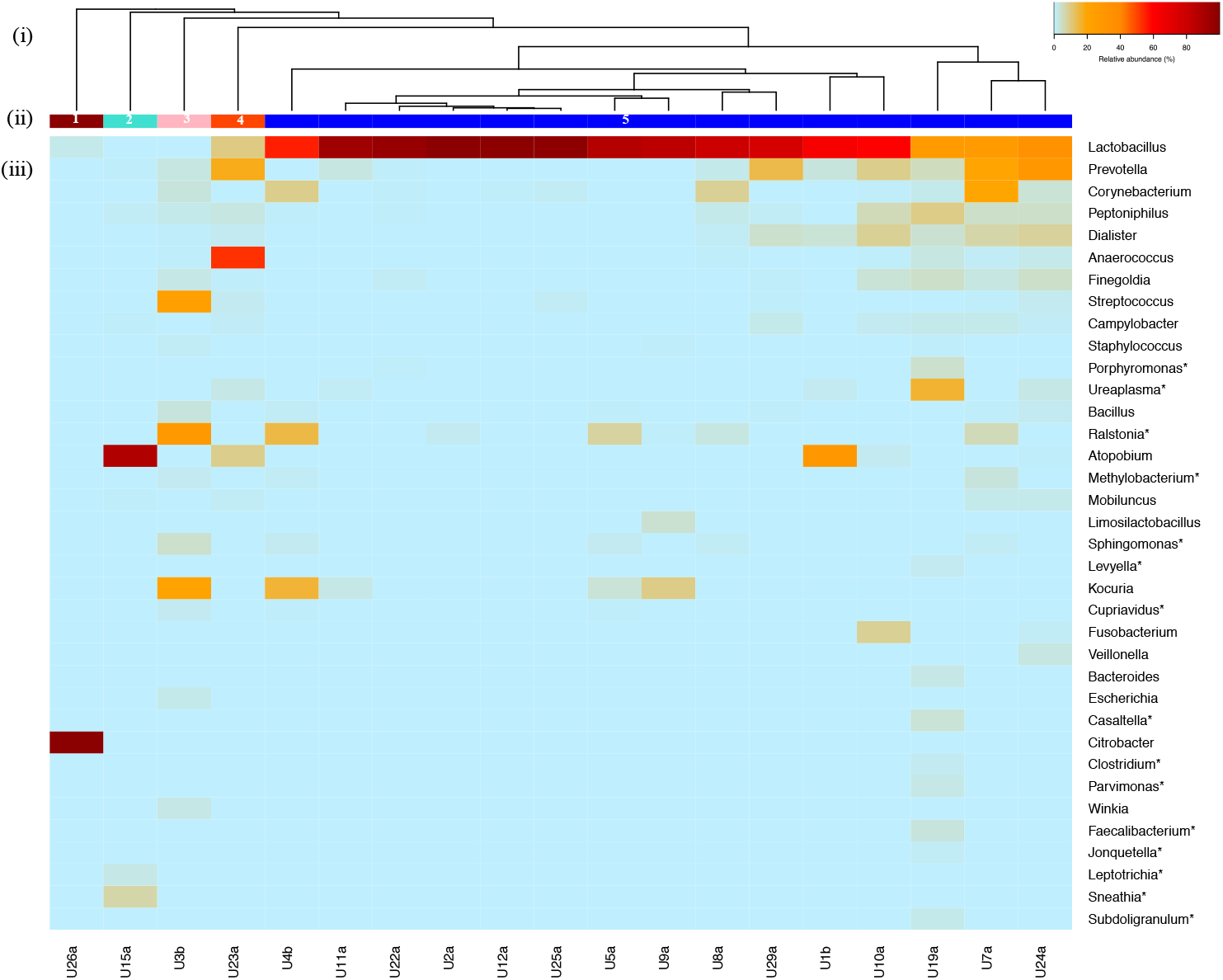
Genus-level community structure types of healthy FUM by community amplicon sequencing. (i) Hierarchical clustering of Bray-Curtis dissimilarity distance matrices on the relative proportions of reads for each OTU within individual urine samples. (ii) Bars below dendrogram denote community structure types. (iii) Heatmap of relative abundances of bacterial genera within each urinary microbiota. Only genera that are at least 1% abundant in at least one sample are shown in order of decreasing prevalence (from top to bottom). Asterisk denotes detection only by community amplicon sequencing and not by culturomics.

**Figure S4.**
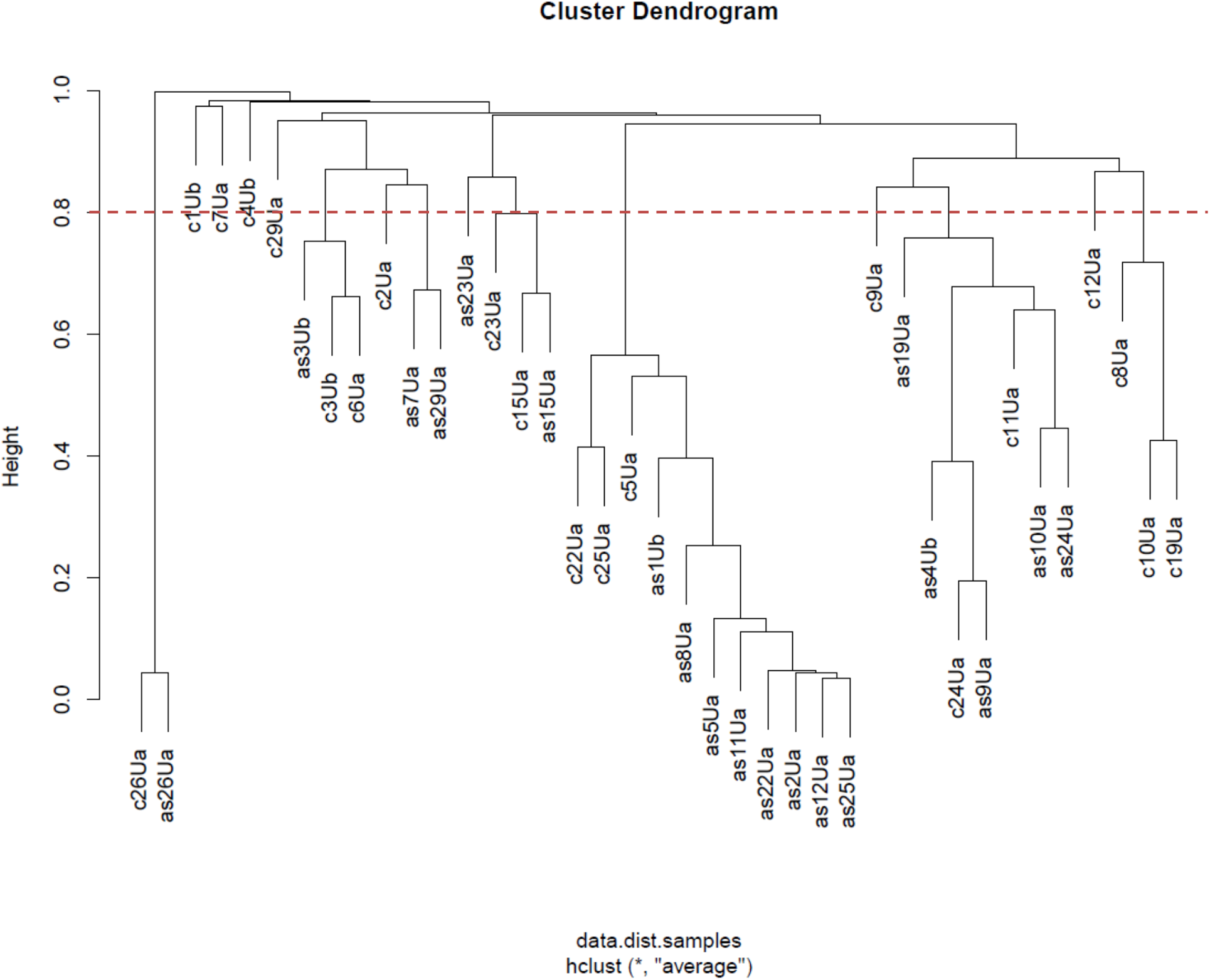
Dendrogram representing species FUM hierarchical clustering including 19 samples characterized by culturomics and amplicon sequencing. Hierarchical clustering was based on Bray-Curtis dissimilarity distance matrices, for which all species detected were included. A cutoff value of 0.8 was used to define the clusters (dashed orange line).

**Figure S5.**
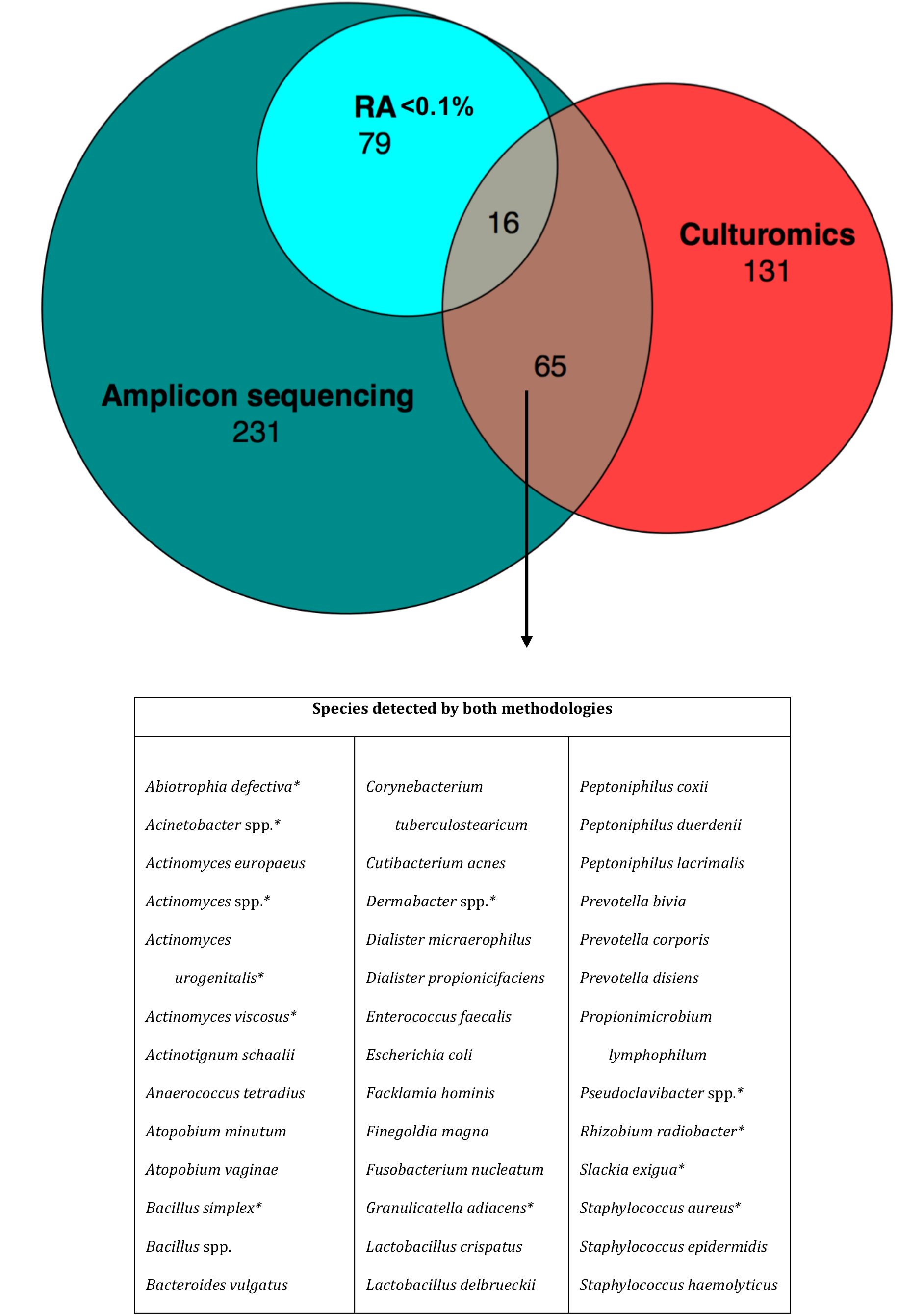
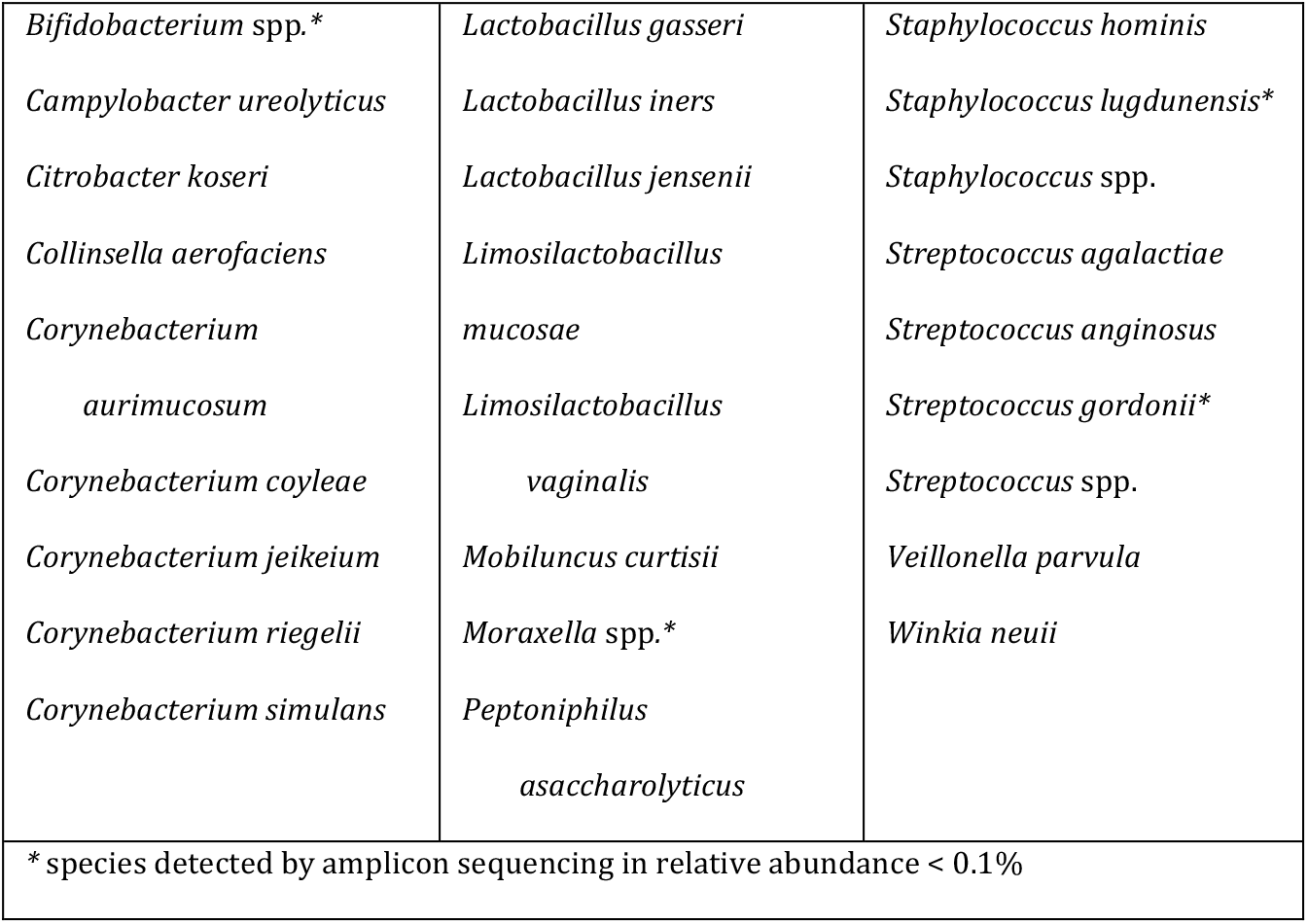
Venn-Euler diagram showing the number of species (N = 297) detected by culturomics and/or amplicon sequencing. The size of the circles and intersections is proportional to the number of species detected. Species detected by both methodologies are listed in alphabetical order. (RA, relative abundance)

